# Contrasting autoimmune and treatment effects reveals baseline set points of immune toxicity following checkpoint inhibitor treatment

**DOI:** 10.1101/2022.06.05.494592

**Authors:** Chen Zhao, Matthew P. Mulè, Andrew J. Martins, Iago Pinal-Fernandez, Renee N. Donahue, Jinguo Chen, Jeffrey Schlom, James L. Gulley, Andrew Mammen, John S. Tsang, Arun Rajan

## Abstract

Immune checkpoint inhibitors (ICIs) have changed the cancer treatment landscape, but severe immune-related adverse events (irAEs) can be life-threatening or fatal and may prohibit patients from receiving further ICI treatment. While the clinical features of irAEs are well documented, molecular signatures, predictive biomarkers, and mechanisms of impending irAEs are largely unknown. In addition, the markers and mechanisms of ICI-induced antitumor immunity often overlap with those for irAEs. It is thus critical to uncover signatures associated specifically with irAEs but not with antitumor immunity. To identify circulating immune cell states associated with irAEs, we applied multimodal single cell analysis (CITE-seq) to simultaneously measure the transcriptome and surface proteins from peripheral blood mononuclear cells (PBMCs) collected before and after treatment with an anti-PD-L1 antibody (avelumab) in patients with thymic cancers (thymic epithelial tumors). All patients had an antitumor response, yet a subset developed muscle autoimmunity (myositis), a potentially life-threatening irAE. Mixed-effect modeling disentangled cell type-specific transcriptional states associated with ICI treatment responses from those of irAEs to identify temporally stable pre-treatment immune set points associated with irAEs only. These pre-treatment baseline signatures of irAE developed post-avelumab irAEs reflect correlated transcriptional states of multiple innate and adaptive immune cell populations, including elevation of metabolic genes downstream of mTOR signaling in T-cell subsets. Together these findings suggest putative pre-treatment biomarkers for irAEs following ICI therapy in thymic cancer patients and raise the prospect of therapeutically dampening autoimmunity while sparing antitumor activity in cancer patients treated with ICIs. Together, pre-treatment biomarkers and interventional therapeutics could help mitigate treatment discontinuation and improve clinical outcomes.

## Introduction

Immune checkpoint inhibitors (ICIs) have demonstrated durable benefit and improved survival in a subset of patients with advanced cancers ^1^. However, this therapeutic benefit comes with a risk of immune-related adverse events (irAEs), common side effects of checkpoint inhibitor therapy ranging in frequency between around 50–90% depending on the type of cancer and checkpoint inhibitor.^2-4^ These autoimmune reactions can be life-threatening, and can affect almost any organ system, with the most common symptoms being rash, pruritus, fatigue, and diarrhea.^4^ While a majority of irAEs can be safely managed by discontinuing ICI treatments and/or giving low-dose steroids, some patients require high-dose steroids or anti-cytokine agents^5,6^,which can decrease the antitumor effect of ICIs. Patients experiencing mild or moderate irAEs can be re-challenged with ICIs under close monitoring^7^; however, the risk of developing a subsequently fatal irAE often precludes continuation of treatment for patients developing severe autoimmunity. There is thus an urgent need for unbiased identification of molecular phenotypes associated with irAE risk to help inform potential biomarkers and treatment strategies to dampen autoimmune effects while sparing antitumor immunity.^8^

Factors contributing to autoimmunity versus antitumor immunity in patients receiving ICIs remain unclear.^9^ Immune inhibitory receptors targeted by these drugs play essential roles in maintaining self-tolerance, as documented in patients with germline mutations affecting these receptors and in transgenic mouse models lacking immune checkpoint inhibitory receptors.^10-12^ Inhibition of negative feedback on immune activation by ICIs may thus cause autoimmune reactions in cancer patients by exacerbating pre-existing clinical or subclinical autoimmunity by increasing the probability of loss of immune tolerance.^13^ IrAE rates are higher in patients treated with dual ICIs, yet single-agent ICI treatment is sufficient to cause autoimmunity in around 20% of patients.^14,15^ Individual variations in baseline (i.e., pre-treatment) immune status (or “set points”^16-18^) may, for example, provide different levels of buffering (or pre-disposition) to develop adverse events. For example, a single “hit” to certain regulatory pathways might be sufficient to cause pathology in some (e.g., those with less buffering capacity^19^) but not all patients. Identifying baseline pre-treatment molecular signatures and states associated with irAE outcomes could uncover biomarkers of immune toxicity with which to select patients for treatment and inform potential treatment interventions. A recent study shed light on the local reaction of T cells at the onset of irAE-related colitis, finding cycling T cells and alterations in T regulatory cells associated with irAEs.^20^ Another report suggested baseline activated CD4 memory T-cell abundance could serve as a biomarker of post-treatment severe irAEs.^21^ Despite these advances, previous unbiased systems-level analyses often used profiling approaches that have limited cellular resolution. Furthermore, statistical assessment of *differences* between signatures associated with irAEs and antitumor responses is lacking, but is critical for understanding the delicate interplay and shared mechanisms between ICI-induced autoimmunity and antitumor immunity. Biomarkers that specifically mark irAEs but not antitumor immunity could help in the development of interventional strategies with minimal impact on the efficacy of ICI therapies.

### Study Design

We set out to contrast treatment-associated and irAE-associated immune system states by profiling peripheral immune cells of patients with metastatic thymic cancer before (baseline) and after administration of the anti-PD-L1 antibody avelumab (at the time or irAE development, or its equivalent in patients not developing irAE). We chose to study irAEs in thymic cancer for the disease’s stable tumor cell-intrinsic property (low tumor mutation burden), good response to ICIs, and high incidence of irAEs.^22,23^ Prior studies in other cancers using cytometry^24^ and single-cell RNA sequencing^25^ investigated responses to ICIs,^25,26^ yet the transcriptional state and phenotype of well-resolved immune cell populations before and after treatment are understudied, particularly involving contrasting treatment- and irAE-associated effects. We addressed these gaps by using CITE-seq (Cellular Indexing of Transcriptomes and Epitopes by Sequencing), a multimodal technique combining surface protein phenotyping and transcriptome profiling simultaneously in single cells, followed by mixed-effects modeling to identify cell subset-specific signatures associated with the development of irAE but not with clinical outcome. Our CITE-seq antibody panel targeted 82 surface proteins and included 4 isotype controls, as previously described.^16^ Nine patients were chosen for CITE-seq analysis; all had clinically similar antitumor activities based on RECIST (Response Evaluation Criteria in Solid Tumors). While no patients had clinically observable autoimmune disease at baseline, five individuals developed myositis after an average of two doses of avelumab (Fig. S1a Supplementary Table 4; see Methods). Paired peripheral blood mononuclear cell (PBMC) samples from baseline (pre-treatment) and at the onset of irAEs post-avelumab (two cycles post-avelumab for the non-irAE group) were used for analysis. Our dataset included more than 190,000 cells from 18 PBMC samples, with two timepoints per patient (Fig. 1a) and a median of 10,804 cells per sample (Fig. S1b,c).

**Figure 1:**
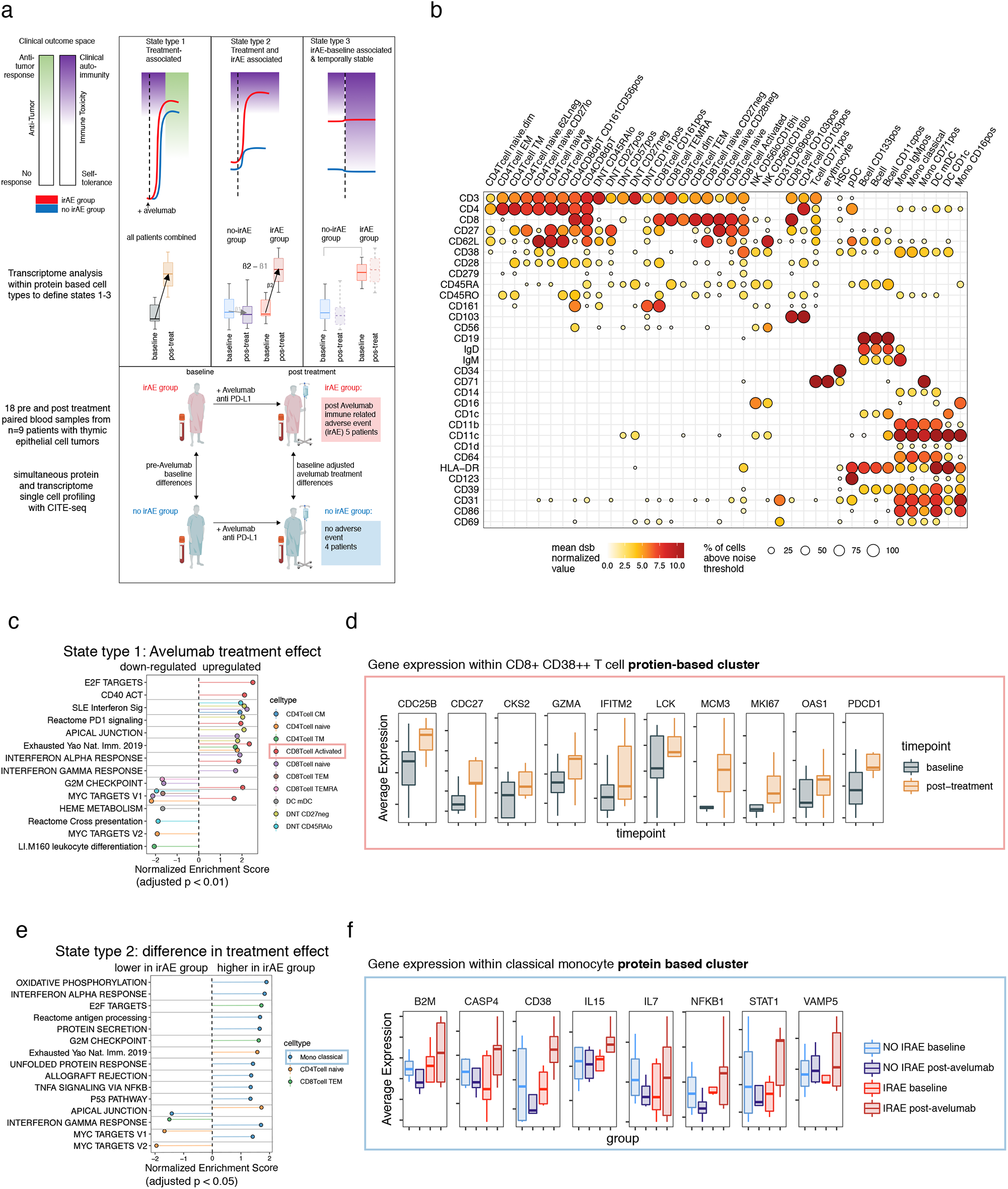
Multimodal single-cell analysis deconvolves transcriptome states associated with irAEs and ICI treatment within protein immune phenotypes. **a**. Top: hypothesized schematic illustrating different types of immune cell states and how those measured parameters reflect the clinical phenotypes: the antitumor response effects (white-blue) and autoimmune toxicity (green-red). Red lines represent the group of patients developing irAEs after treatment and blue represents those without irAEs. Cell states from left to right: State type 1: perturbed by treatment across all patients which can be associated with antitumor effects but also with autoimmune toxicity; State type 2: increased post-treatment in the irAE group (these could reflect a higher fold change in the irAE group or oppositely regulated states); State type 3: baseline differences in the irAE group exhibiting temporal stability over the course of treatment, which is associated with irAEs but not treatment effects. Middle: transcriptome comparisons carried out within protein clusters to identify different cell states corresponding to the states above. Bottom: study scheme devised to define cell state differences above: eighteen PBMC samples from nine patients with thymic cancers were profiled at baseline and post-avelumab treatment; five of them developed an irAE (myositis) post-treatment and the other four patients did not develop an irAE. PBMC samples collected before treatment (baseline) and post-avelumab (at the onset of irAE and matched time points in the non-irAE group) were profiled using CITE-seq with a panel of 82 antibodies. **b**. CITE-seq surface protein expression map of PBMCs. Circle color is the mean dsb ‘denoised’ and normalized protein level; the scale of dsb values can be interpreted as the number of standard deviations above background noise. Circle size is the percentage of cells in the cluster that express the protein above the expression-positive cutoff of 3.5. **c**. State type 1: treatment effects : gene set enrichment based on genes ranked by the pre- vs. post-avelumab treatment effect from donor-weighted pseudobulk model. **d**. Selected genes from the CD38^++^CD8^+^ effector T-cell cluster in leading-edge genes of the enriched pathways shown in c. **e**. State type 2: difference in treatment effects between the irAE and non-irAE group: gene set enrichment. **f**. Selected genes from the classical monocyte cluster with a treatment effect upregulated in the irAE group – genes include those with oppositely regulated directions (CD38 mRNA) or genes only perturbed in the irAE group (IL15).

### High dimensional protein-based immune cell phenotyping

Defining cell clusters and subsets with surface protein alone allowed us to identify cell type based on well-studied surface markers (Fig. S1d), thereby separating transcriptome measurements from cell type identity. This facilitated improved interpretation of transcriptome differences between outcome groups within cell clusters that were defined with statistically independent (protein) information from transcriptome data (Fig. 1a). We clustered cells using spectral clustering based on the denoised expression level of 82 surface proteins. This procedure identified 43 cell clusters spanning major cell lineages, including subsets of B cells, monocytes, dendritic cells (DCs), natural killer cells, and T cells (Fig. 1b, Fig. S1b). The substantial number of antibodies in our panel for marking T-lymphocyte phenotypes and cell states revealed significant heterogeneity within CD8^+^ and double negative (CD3^+^CD4^−^CD8^−^) T-cell subsets, with most of these clusters/phenotypes detected across donors (Fig. S1c).

### Statistical modeling of avelumab treatment and toxicity effects between patient groups

Our analysis approach focused on defining cell states associated with irAEs decoupled from treatment response within immune cell types defined by protein. To this end, we applied statistical contrasts to identify changes of cell functional states due to three major effects: 1) <u>ICI treatment effects</u>–pre- vs. post-avelumab treatment effects shared across all subjects; 2) <u>ICI-associated irAE effects</u>–the difference in pre- vs. post-treatment effect between irAE and non-irAE groups; and 3) <u>baseline effects</u>–differences in cell state prior to avelumab treatment between groups (analogously shown in Fig. 1a). By subtracting treatment effects (both effects 1 and 2) from baseline differences, we further focused on baseline cell states associated with impending irAEs exhibiting temporal stability over the course of treatment; these cell states were thus uncoupled from ICI response effects. To accommodate our experimental design containing patients with repeated measurements nested in groups, we used weighted mixed-effects models at the single-cell level and on pseudo-bulk data aggregated within each cluster to model variations across donors over time and between outcome groups (see Methods). We tested enrichment of a pre-specified list of gene modules based on our hypothesis of pathways that could tune immune states related to irAEs and response (see Methods, Supplementary Table 3), in addition to carrying out unbiased analysis of 50 MSigDB Hallmark pathways.

### Defining ICI treatment effects

We first assessed cell type-specific avelumab treatment effects (Fig. 1a-State type 1) by identifying, within each cell cluster above, the transcriptional differences between post- vs. pre-treatment across patients (detailed results are provided in Supplementary Table 1). Transcriptional signatures of T-cell activation, interferon pathways, PD-1 signaling and T-cell exhaustion were elevated within multiple T-cell subsets (Fig. 1c, Supplementary Table 1). Activated CD38^++^ CD8+ T cells and naïve CD8^+^ T cells had the highest number of enriched pathways. In the CD38^++^ subset, upregulation of T-cell activation and cell cycle signals included genes *CDC25B, CDC27, MCM3*, and *CSK2* (Fig. 1d). Upregulation of *GZMA, OAS1, IFITM2, LCK, MKI67*, and *PDCD1* in this subset driven by elevated activation protein CD38 was consistent with a proliferating effector phenotype (Fig. 1d). Interestingly, cell cycle states were enriched in the opposite direction (downregulated post-avelumab) in several other T-cell subsets including CD8^+^ T_EMRA_ and CD8^+^ naïve T cells (Fig. 1c). Genes driving pathway enrichments tended to be mutually exclusive among different cell clusters/subsets (as defined by the Jaccard similarity of the “leading edge” genes for each gene set/pathway; see Methods) (Fig. S2a and Supplementary Table 1). The presence of cell cycle signatures in CD38^++^ effector T cells post-avelumab treatment is reminiscent of phenotypes revealed in prior studies, e.g., those focusing on the dynamics of T-cell changes during ICI treatment^26,27^ and a study of ICI-induced colitis.^20^ Together, our and others’ observations suggest that a proliferation signature of peripheral CD8^+^ effector T cells is coupled to ICI treatment responses.

### Defining ICI treatment effects unique to irAEs

While ICI may trigger qualitatively similar responses in different patients, the quantitative extent of these responses may differ between irAE and non-irAE patients. To evaluate this possibility, we identified treatment response-associated irAE effects by comparing the difference in post-treatment vs. baseline fold changes between the irAE and non-irAE groups (Fig. 1a, State type 2); this analysis identified 35 enrichments involving 14 cell types (Supplementary Table 1). Within classical monocytes in particular, interferon transcriptional signatures were highly enriched (Fig. 1e) and leading-edge genes in these pathway enrichments tended to be mutually exclusive (Fig. S2b). *IL15* and interferon-simulated genes (ISGs) *B2M*, CD38 and *STAT1* were oppositely regulated between groups from baseline to post-treatment (upregulated in the irAE group and downregulated in the non irAE group). These further suggested that interferon-responsive monocytes were activated in the irAE group after treatment with avelumab (Fig. 1f). This IFN response in monocytes that is associated with irAEs may phenocopy the elevated interferon response states seen in inflammatory diseases such as lupus,^28^ and has also been observed in patients with myositis.^29^ In addition, cell cycle/proliferation signals were enriched post-treatment in the irAE compared to the non-irAE group in CD8 effector memory T cells (Fig. 1e).

### A baseline metabolic transcriptional signature is associated with post-treatment irAEs independent of treatment effects

We next searched for signatures of baseline (prior to treatment) immune states in patients who developed irAEs post-treatment. We focused on signatures uncoupled from avelumab treatment effects. We first identified baseline states associated with post-treatment irAEs, then subtracted enrichment signals associated with avelumab treatment (see Methods). This procedure resulted in a map of temporally stable cell type-specific signatures, or “set points”, associated with the post-treatment development of irAEs independent of treatment effects (i.e., those defined by Fig. 1c-f, State type 1 and 2 in Fig. 1a). Inflammatory and metabolic signatures including mTOR and TNFα pathway genes were enriched within multiple cell subsets (Fig. 2a). DCs have known roles in modulating autoimmune and antitumor responses,^30^ and both CD1c^high^ and CD1c^low^ DCs in the irAE group appeared to have an elevated inflammatory signature at baseline, e.g., TGFB, TNFA, and inflammatory response pathway enrichments. They also displayed potential enhanced tissue migratory capacity given the enrichment of epithelial-to-mesenchymal transition genes, including *CD44, VIM, VCAN, THBS1*, and *SDC4* (Fig. 2a, Supplementary Table 1). These primed DC subsets were also elevated for several metabolic-related transcript differences, for example, involving mTORC1 signaling, hypoxia, and cholesterol homeostasis. Some of these inflammatory and metabolic signatures are also shared by CD8^+^ T cells, in particular memory cells re-expressing CD45RA (CD8^+^ T_EMRA_), which displayed elevated TNF signaling, hypoxia, cholesterol homeostasis, and mTOR signatures in the irAE group (Fig. 2a). By design, these phenotypes we uncovered were not enriched in the irAE group after avelumab treatment, where innate immunity and inflammatory signatures such as those associated with IFNs were more specific to CD14^+^ classical monocytes (Figs. 1e,f).

**Figure 2:**
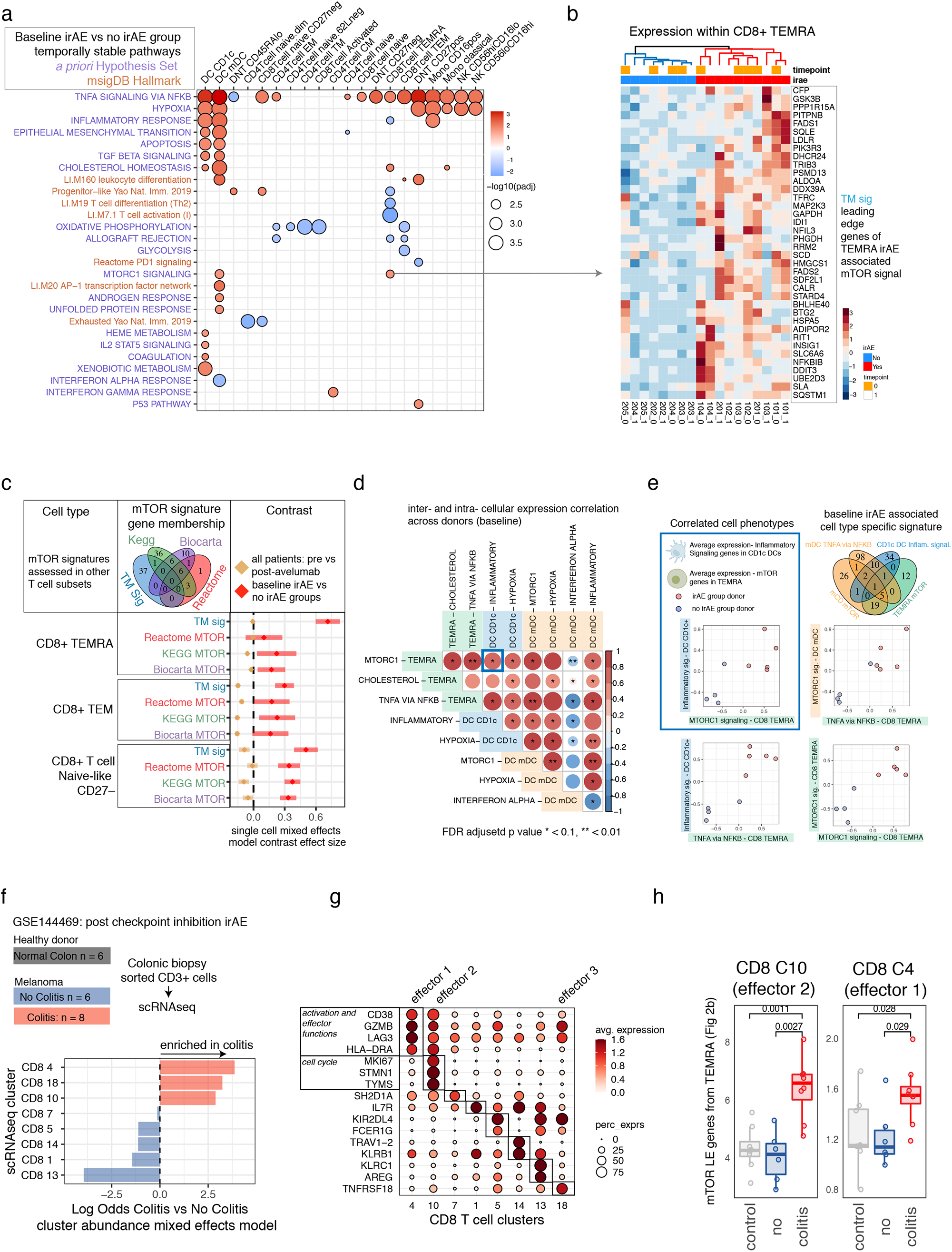
baseline immune set points associated with post-treatment ICI toxicity. **a**. Baseline gene set enrichment map based on genes ranked by weighted pseudobulk model effect size within each cluster comparing patients with eventual irAEs to those without irAEs. Red and blue indicate pathways positively and negatively enriched respectively in patients before avelumab treatment who developed an irAE after receiving avelumab. The pathway names (y-axis) highlighted in blue are from the MSigDB hallmark collection, and pathways in dark orange are curated gene sets based on a pre-study defined hypothesis (Supplementary Table 3). **b**. Average single cell expression of leading-edge genes from the Hallmark mTOR pathway baseline enrichment associated with the irAE group within the T_EMRA_ cell cluster. Samples from both time points are ordered according to hierarchical clustering with complete linkage. **c**. The coefficient corresponding to baseline irAE vs. no-irAE (red) and the fold change across donors (tan) from a single-cell mixed-effects model of other mTOR signatures across CD8 T-cell subsets. Tan lines with a coefficient effect size near 0 are temporally stable; pathways with an effect size above 0 are associated with irAEs; error bars are 95% confidence intervals of the contrast applied to mixed-model fits. **d**. Sample-level baseline pseudobulk expression correlations of temporally stable baseline cell states associated with development of irAEs after avelumab treatment. Each box represents the Pearson correlation coefficient (two-sided) of donor pseudobulk data with the FDR adjusted p-value (FDR adjustment across all temporally stable baseline enrichments, subset of states shown in correlation matrix) shown with asterisks. **e**. Scatterplots of average donor expression for selected inter- and intracellular correlations shown in 2d. **f**. Data from Luoma *et. al*. 2020 (GSE144469) which profiled patients treated with combined checkpoint inhibitors who went on to have suspected colitis that was either confirmed on biopsy with overt colitis (red) vs. no evidence of colitis (blue) and healthy colon biopsies (grey) with CD3^+^ cells FACS sorted followed by single-cell RNA-seq. Bottom shows association testing using an aggregated binomial generalized linear mixed model of the association of cells from each cluster with the colitis vs. no colitis groups. **g**. Expression of selected differentially expressed genes for each cluster of colonic T cells from a one cluster vs. all ROC test (Seurat). **h**. The average colitis T-cell expression of the mTOR-LE gene signature within effector CD8 T-cell clusters 10 and 4 across donors.

### Correlated cell-state phenotypes underlie baseline set point signatures of irAEs

Given the critical role of the mTOR pathway in tumorigenesis^31^ and autoimmunity,^32^ we further examined the genes driving mTOR pathway enrichment within CD8 T_EMRA_ cells (Fig. 2a). These leading-edge mTOR genes (mTOR-LE) naturally clustered in all samples, independent of time points, into two clusters segregated by irAE status (Fig. 2b). The independence from time points confirmed that our procedure identified temporally stable enrichment signals that were stable between the pre- and post-treatment time points. Expression of mTOR-LE genes was elevated in thymic cancer patients compared to both the non-irAE group and healthy donors (n=20) assessed with the same CITE-seq panel within gated CD8 T_EMRA_ cells (Fig. S3a-c). mTOR-LE genes such as *SLC2A1, GAPDH, FADS1, FADS2, LDLR*, and *ADIPOR2* (Fig. 2a) suggested this enrichment may reflect a metabolic state downstream of mTOR, since these genes are involved in glucose and lipid metabolism. Therefore, we further tested 6 distinct mTOR signatures covering different aspects of the pathway from public databases (Fig. S3e) by repeated differential expression models using a k-cell permutation approach (see Methods). All mTOR signals as well as the TNF pathway were consistently enriched in the irAE group in CD8 T_EMRA_, (Fig. S3e), suggesting the mTOR-LE signal may have reflected an immune state controlled in part by upstream mTOR signaling. We next wondered if this state could be shared by other cell types. Using a more sensitive model accounting for variation at the single cell level and modeling expression of the 6 mTOR pathways defined above (see Methods) revealed mTOR signatures elevated at baseline in the irAE group within double-negative (DNT: CD4^−^CD8^−^), double-positive, and CD8^+^ subsets including CD8^+^ T_EMRA_, T_EM_, CD27^−^ naïve-like cells (Fig. 2c and Supplementary Table 2), while CD4 subsets were not enriched for mTOR pathways (Supplementary Table 2). Consistent with temporal stability of the original CD8 T_EMRA_ mTOR-LE signature, baseline elevation in the irAE group was uncoupled from treatment effects in these subsets (Fig. 2a, tan estimate and errors near 0). Notably, mTOR signatures were not upregulated in the irAE group within the activated CD8^+^ CD38^++^ T-cell cluster which had a post-treatment phenotype overlapping with antitumor responses^33^ (see above Fig. 1c-d). To investigate molecular identity of these protein-based cell types independent of the irAE group differences, we integrated healthy donors’ and thymic cancer patients’ CITE-seq data into a joint CD8 T-cell map (Fig. S4a-d). CD8 T_EMRA_ were distributed across multiple clusters including cluster 1, which was enriched for mTOR-LE genes and expressed genes controlling terminal effector fate *HOPX*,^34^ *GZMH*,^35^ and the transcription factor *ZEB2*^36^ (Fig. S4c).

To further characterize shared information between distinct baseline irAE-associated enrichment signals, we correlated baseline expression of enriched pathway leading-edge genes across subjects both within and between cell types (Fig. 2d, Fig. S2c). Supporting our hypothesis that CD8 T_EMRA_ mTOR captured a more global irAE-associated metabolic state, as reflected by mTOR’s elevation across different effector and naive CD8 subsets in the irAE group (Fig. 2c), CD8 T_EMRA_ mTOR-LE expression correlated with other irAE-associated metabolic and inflammatory signaling pathways across subjects (top row of Fig. 2d). For example, the CD1c DC inflammatory signaling and T_EMRA_ mTOR phenotypes were correlated across donors (inset in blue in Fig. 2e). The level of the T_EMRA_ mTOR signal was correlated with T_EMRA_ TNF signaling (Fig. 2e) and this TNF signal was associated with the level of innate-cell mTOR and inflammatory signaling (Fig. 2d, e). The irAE baseline innate inflammatory and metabolic phenotype appeared distinct from states related to interferon tone, as mDC interferon signaling was negatively enriched in the irAE group and negatively correlated with metabolic and inflammatory states across subsets. Together, these correlated cell phenotypes suggest stable inter- and intracellular rewiring of inflammatory and metabolic states comprising a shared immune set point of patients primed toward development of autoimmunity after treatment with avelumab.

### Assessing irAE-associated T-cell signature in tissue-localized T cells associated with ICI-induced colitis

The circulating CD8^+^ T-cell set point signature we identified may phenotypically overlap with those found in tissues associated with adverse immune reactions. We investigated this further by assessing our CD8^+^ T-cell signatures in single-cell RNA sequencing data obtained from a published study of CD8^+^ T cells isolated from colonoscopy biopsies of healthy donors (n=8) and patients with melanoma with (n=8) and without (n=6) active ICI-induced colitis^20^ (Fig. 2f, Fig. S5a). We focused on eight CD8^+^ T-cell clusters defined by unsupervised clustering; three of these CD8 clusters were specific to the CD8^+^ T cells isolated from colitis lesions (Fig. 2F), all of which had an effector phenotype based on mRNA expression (Fig. 2g). We further examined cluster 4 (“effector 1”) and cluster 10 (“effector 2”) as these had sufficient numbers of cells after aggregation across subjects (see Methods). Within these T-cell clusters, donors with colitis had higher relative expression of the mTOR-LE gene signature compared to T cells from either healthy donors or ICI-treated melanoma patients without colitis (Fig. 2h). Interestingly, *BHLHE40*, a member of our mTOR-LE signature, is the gene with the most significant differential expression between ICI-induced colitis vs. non-colitis in cluster 10 (effector cluster 2). This gene is a crucial regulator of cytokine production associated with autoimmune responses,^37^ which is consistent with the notion that the mTOR-LE gene signature reflects a “poised” metabolic/inflammatory phenotype.

## Discussion

In this work we identify a set of highly interpretable multimodal molecular states associated with ICI response and adverse events. We found multiple cell functional states linked to ICI response were also likely coupled to those involved in irAEs. However, patients with post-treatment irAEs shared a common baseline immune set point, reflecting elevated inflammatory tone and metabolic differences across the innate and adaptive immune systems. Our analysis suggested that this baseline set point may be tuned by common upstream regulators such as mTOR, which is known to regulate metabolic states such as hypoxia.^38^ Given the role of mTOR inhibitors as both antitumor agents and suppressants of autoimmunity, our results provide the rationale for evaluating the concurrent use of mTOR inhibitors with ICI in an attempt to diminish the risk of developing irAEs while preserving the antitumor effects of ICI. Intriguingly, a case report of a renal allograft patient with melanoma treated with an ICI and an mTOR inhibitor found the antitumor effect could be preserved while the autoimmune toxicity could be limited.^39^ However, further studies are needed to confirm these observations.

Our study has several limitations. Multimodal single-cell profiling of more than 190,000 cells created a high-resolution map of cell states, but the number of patients included in our study is limited and, due to experimental constraint at the time of this experiment, samples were split across two batches. We have previously found staining batch has limited impact on technical effects in CITE-seq data ^40 16^. However, assessment of batch effects across sample groups was limited due to the small sample size confounding in this dataset. Future single-cell analysis of irAEs could include additional subjects to assess the generalizability of the molecular states we derived herein. The generalizability of these findings to other types of cancers and ICIs could also be assessed in future work, although we did find overlap in signals within tissue autoimmunity from melanoma patients treated with different ICIs. Finally, it will be interesting to link differences in immune cell states detected from blood with those at the tissue level from sites of involvement by the irAEs. While these clones would be difficult to trace in humans, lineage tracing mouse model systems could be informative in studying the origins of these cells. Together, our dataset, analysis, and curated results can serve as both a framework and a rich source of hypothesis-generating data to inform future precision immunotherapy research on biomarkers and treatment strategies for irAEs.

## Author contributions

C.Z., A.R., and J.S.T. conceived and designed the study; M.P.M., C.Z., A.J.M., and J.S.T. designed data analysis strategies; M.P.M. and A.J.M. performed data analysis with help from C.Z.; C.Z., A.R., J.L.G., I.P.F., and A.M. performed the clinical study; J.S. and R.D. collected samples and analyzed clinical information; M.P.M., A.J.M., C.Z., and J.S.T. designed CITE-seq experiments; M.P.M., A.J.M., J.C., and I.P.F. generated CITE-seq data; C.Z., M.P.M., and J.S.T. wrote the manuscript with contributions from A.J.M.; C.Z. and J.S.T. supervised the study; all authors approved the manuscript.

## Acknowledgements

C.Z., R.D., J.S., J.L.G., and A.R. are supported by the Intramural Research Programs (IRP) of NCI (Center for Cancer Research); M.P.M., A.J.M., J.C., and J.S.T. are supported by the IRP of the NIAID; I.P.F. and A.M. are supported by the IRP of the NIAMS; this research was supported by a Cooperative Research and Development Agreement between the National Cancer Institute and EMD Serono, Billerica, MA, USA (CrossRef Funder ID: 10.13039/100004755), as part of an alliance between the healthcare business of Merck KGaA, Darmstadt, Germany and Pfizer. The healthcare business of Merck KGaA, Darmstadt, Germany (CrossRef Funder ID: 10.13039/100009945) and Pfizer reviewed this manuscript for medical accuracy only before journal submission. The authors are fully responsible for the content of this manuscript, and the views and opinions described in the publication reflect solely those of the authors. We thank members of the Tsang Lab and the NIH Center for Human Immunology (CHI) for inputs and discussions. We thank Can Liu for helping with the sequencing process. We thank Drs. Pamela Schwartzberg and Ronald N. Germain for critical feedback on the project. We thank Shannon Swift, Susan Sansone, and Madison Ballman for clinical support and provision of samples. Illustrations were created with BioRender.com.

## Methods

### Clinical/sample collection

Patients with advanced thymic cancers were enrolled in clinical trial NCT03076554 approved by the NCI’s Institutional Review Board and received avelumab, an anti-PD-L1 antibody, every two weeks. PBMC samples were collected before starting therapy, at the end of every treatment cycle, and at the onset of irAEs. No patients had any history of autoimmune disease prior to treatment. Patient characteristics are listed in Supplementary Table 4.

### Multiplexed CITE-seq single-cell transcriptome and protein profiling

Cells were thawed in RPMI with 10% FBS and washed and stained in 1xPBS with 0.04% BSA. CITE-seq was performed as previously described in Kotliarov et al. using the same antibody panel. Donor cells were stained with sample barcoding antibodies,^41^ washed, and pooled into a single tube; two staining batches were used to accommodate a greater number of samples than available barcode antibodies. Although a single batch design was planned using lipid indexing ^42^, a subset of samples had red cells noticeable in the PBMC prep; we therefore used HTO staining in 2 batches due to unknown effect of residual RBC membranes on LMO staining (Supplementary Table 4).Pooled cells were stained with a concentrated optimized panel of 86 antibodies (including 4 isotype controls; anti-mouse (rather than anti-human) CD206 was incorrectly included in the panel and not considered in the analysis). The stained cell pool was then washed and prepared according to the 10X Genomics cells partitioned across eight lanes of the 10X Genomics chromium microfluidic instrument per staining batch. Sequencing libraries were prepared using the 10X Genomics 3’ assay with version 3 reagents. Antibody-derived tag (ADT) libraries from sample barcode antibodies and surface phenotyping antibodies were prepared according to the publicly available protocol on cite-seq.com. Sequencing was performed on an Illumina NovaSeq system.

### Normalizing and denoising CITE-seq protein levels and protein-based clustering

After sample demultiplexing and doublet removal based on sample barcoding antibodies, CITE-seq surface protein data were normalized and denoised using dsb^43^ to correct protein-specific background noise using ADT reads in empty droplets and correct technical cell-to-cell variations using isotype controls/models fitted to each cell. Default dsb algorithm parameters were used (denoise.counts = TRUE, use.isotype.controls = TRUE). The normalized values were then batch corrected using limma. Single cells were clustered using a Euclidean distance matrix formed from the normalized protein values as input to spectral clustering using Seurat version 3.1.5.^44^ A total of 44 cell clusters were annotated based on protein expression.

### Analysis of aggregated transcriptome data within protein-based clusters

Gene expression counts were aggregated into a pseudo-bulk library within each protein-based cluster by adding counts for each sample x cell type into a summed count matrix, and cell types without representation (e.g., donor-specific clusters) were excluded from analysis. The aggregated counts for the n=18 samples across each cell type were normalized using the trimmed means of M values method^45^ and genes were retained which had a pooled count per million above 3 across sufficient samples based the edgeR *filterByExprs* function. Filtering genes in a cell type-specific manner removed genes from analysis unexpressed by a given cell type (e.g., genes specific to a different lineage) and ensured assumptions of the model to derive precision weights^46^ used to account for variations in sample quality/library size were met, i.e., the log count per million vs. fitted residual square root standard deviation had the expected monotonically decreasing trend within each cell type (see below).

### Estimating subject and group-level effects within protein-based clusters

Target estimates of statistical analysis were treatment and group-level transcriptional effects within the protein-based clusters defined above. Models were fitted to single-cell and aggregated data (see below). To assess these effects, we contrasted fitted values of fixed and mixed effects linear models of gene expression within protein clusters. A contrast matrix *L* was constructed with a single combined factor variable group.time corresponding to irAE outcome group and time point relative to treatment with levels 1 = irAE baseline, 2 = irAE post-avelumab, 3 = no irAE baseline, 4 = no irAE post-avelumab (columns, below). The matrix was used to make the following comparisons (rows) based on fitted model values (see below) 1) <u>ICI treatment effects</u>–across all subjects, 2) <u>ICI-associated irAE effects</u>–the fold change difference between groups and 3) <u>baseline effects</u>–the baseline difference between the irAE and non-irAE group.

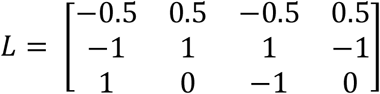

### Mixed effects models on aggregated data to estimate treatment effects across donors and fold change differences between groups

Estimation of avelumab treatment effect across all donors and the difference in treatment effects between irAE groups was modeled with a mixed-effects model including a varying effect for subject ID to model variation in baseline expression. Models were implemented with the variancePartition package ‘dream’ method^47^ to fit models with precision weights in a mixed effects model using lme4.^48^ For each gene we applied the formula f1 = gene ∼ 0 + group.time + (1|subjectID) and fit models using the function ‘dream’. The fitted value for expression y of each gene g corresponds to:

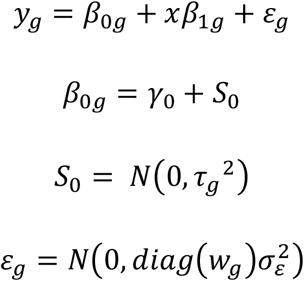

Where *x* represents a combined group.time factor variable defined above, *β*_0_ corresponds to the (1|subjectID) term to model the variation across subjects S around the average *γ*_0_ baseline expression, and errors *ε*_*g*_ are modeled with precision weights *w*_*g*_ calculated using the voom^46^ method. The first two rows of contrast matrix above were applied to estimate 1) the coefficients for the treatment effect across all donors and 2) the difference in fold changes between the irAE and non-irAE groups from mixed-effect model fits.

### Modeling baseline states associated with development of irAE

The third row of the contrast matrix above was used to estimate baseline differences using a fixed-effects model with limma with the function lmFit using voom^46^ precision weights as above in a fixed-effects model. The Empirical Bayes moderated t statistics for each gene comparing the irAE group to the non-irAE group were calculated using the limma^49^ eBayes function. After gene set enrichment (see below “Enrichment testing of hypothesis set and unbiased pathways in model contrasts”), we defined the subset of these baseline states associated with later irAEs which exhibited temporal stability over the course of treatment. The pathways enriched in the irAE group with adjusted p values < 0.01 were further filtered by removing any enrichments evidence of kinetic change (including weak evidence). For each enriched pathway within each cluster, if either the treatment across donors or the irAE-associated treatment effect enrichments (see below “Estimating avelumab treatment effects across donors and between groups”) had adjusted p values of 0.1 or less and were either positively or negatively enriched, these were considered kinetically altered by treatment for the purpose of filtering the baseline signals. These kinetically altered signals were subtracted from the baseline enrichments in a cell type-specific fashion, and the remaining enriched baseline pathways associated with development of post-treatment irAEs were considered temporally stable states.

### Single-cell mixed-effect models

The same formula f1 above (see “Mixed effects models on aggregated data to estimate treatment effects across donors and fold change differences between groups”) was used in a linear mixed model on expression of gene modules within single cells in specific T-cell subsets. The model estimated variation at the single-cell level instead of at the individual donor aggregated level and otherwise corresponds to the same model formula as described above without voom observational weights in the error term, i.e.,

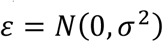

Gene expression of each gene *g* in each cell *i* was normalized log transformed with library size scaling factors using the Seurat function NormalizeData() with *normalization*.*method* = ‘LogNormalize’ to implement the transformation:

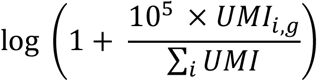

Average expression of gene modules/pathways was then calculated for each module for each single cell and standardized within each protein-based subset by subtracting the mean and dividing by the standard deviation of the average score. Models were fit using the R package lme4 and the treatment effect across donors, and the baseline difference between irAE and no irAE groups was estimated using the contrast matrix *L* above with the emmeans package.^50^ Models were checked for convergence criteria and no models were flagged as having singular fits.

### Enrichment testing of hypothesis set and unbiased pathways in model contrasts

To test enrichment of pathways based on the estimated gene coefficients corresponding to the three effects defined above, we performed gene set enrichment analysis using the fgsea package,^51^ using 250,000 permutations of the ranked gene list to form null distributions for p values; genes were ranked based on the empirical Bayes moderated t-statistic for the baseline comparison of irAE status or with the raw t-statistic for mixed-effect models comparing treatment effects over time. Two gene sets were assessed: first a hypothesis set of modules curated from the Li et al. Blood Transcriptional modules,^52^ MSigDB Hallmark,^53^ Reactome pathways,^54,55^ and curated from literature^56-58^ were tested for enrichment (Supplementary Table 3); the full MSigDB Hallmark pathways were tested independently. The Jaccard similarity of enrichments within cell types was calculated using the geneOverlap^59^ package.

### k-cell permutation profiling of CD8 T-cell signatures and enrichment

To assess the robustness of gene set enrichments, T_EMRA_ cells were manually gated based on dsb normalized CITE-seq protein expression of CD3,CD8,CD45RA, and CD27. The same cells were gated from the previously published data on 20 healthy donors from Kotliarov *et al*., and the average expression of the mTOR-LE genes was compared across thymic cancer irAE groups and healthy donors using a non-parametric Wilcoxon rank test. To account for variability in the number of cells per donor in both manually gated and unsupervised T_EMRA_ clusters, we quantified stability of enrichment to cell sampling variations. We re-ran the pseudobulk baseline differential expression model (as described above “Modeling baseline states associated with development of irAE”) 100 times with libraries constructed from random k-cell samples (without replacement) of 45 cells from each donor. The k value of 45 was chosen as it was the median number of cells in the group (healthy donors) with the lowest number of gated T_EMRA_ across all donors. Pseudobulk libraries were constructed and differential expression testing of the irAE vs. non-irAE groups and healthy donors was carried out as above using limma. Genes within each k sample were tested for enrichment using two complementary methods with highly concordant results. 1) genes were filtered for testing based on the pseudobulk expression profile of the k cell pool with a minimum of 3 counts per million based on the design matrix as above. 2) The same genes as in the original T_EMRA_ cluster were fit with limma regardless of their expression status in the k cell pool. Genes were then ranked by empirical Bayes t statistic comparing irAE vs. non-irAE or irAE vs. healthy donors (not shown and highly concordant with irAE vs. non-irAE, as expected based on the average expression profiles in Supplemental Fig. 3b). Gene set enrichment was assessed using 250,000 permutations of the gene rank list for the null distribution, as described above.

### Integrated analysis of healthy donors and thymic cancer patients

Seurat version 3.1.5^44^ was used to integrate healthy donor and thymic cancer PBMC data using healthy donors as the reference dataset. Regularized Pearson residuals^60^ were used to normalize data for integration with a covariate for subject. Integrated data were clustered using 30 principal components. Differential expression was compared between integrated clusters with an ROC test in Seurat (FindAllMarkers, test.use = ‘roc’). Immune cells from cancer patients often clustered more distinctly; however, this shared state map based only on mRNA helped further define mRNA substates irrespective of the group comparisons described above.

### Analysis of colonic T cells from patients with and without colitis following checkpoint inhibitor treatment from Luoma *et al*. 2020

We reanalyzed the colonic T-cell data from Luoma *et al*. (GSE144469) which included three patient groups: healthy donors, patients treated with ICIs with subsequent colitis (irAE group) and without colitis (non-irAE group). T cells were isolated from the site of the colitis lesion in the irAE group. T-cell single-cell mRNA data were clustered using 30 principal components from 2000 variable genes (FindVariableFeatures with selection.method = ‘vst’) using Seurat version 3. Differential expression of genes between clusters was carried out using a one-cluster vs. all-ROC classifier implemented in Seurat. Cluster association with irAE group status was carried out with an aggregated binomial mixed-effects model to estimate the proportion *p* of cells in each cluster *c* from each subject *S* belonging to each group *g* accounting for within-donor replicated cells (i.e., pseudoreplication) in each cluster. The model formula n/total ∼ IRAE + cluster + IRAE:cluster + (1|subjectid) was fit using lme4 with the glmer function (family = ‘binomial’) and *weights* parameter equal to the total number of cells from each donor, taking the form:

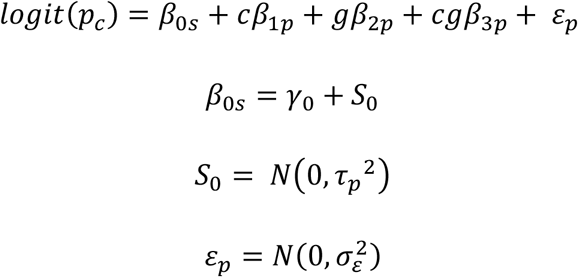

Where p (proportion) was equal to ncells/total cells for each donor, c = cluster ID and g = group ID (irAE vs. no irAE). Data were fit in aggregated format at the level of cells per sample x cell type combination. The fitted marginal means were calculated for irAE group status conditional on cluster ID and back transformed to the log odds space using the emmeans package.

## Code availability

Analysis code and documentation to reproduce this work is available in the repository: https:// https://github.com/niaid/irAE_manuscript

## Data availability

Links to data generated in this study will be made available in the repository: https:// https://github.com/niaid/irAE_manuscript

---

**Supplemental Figure 1.**
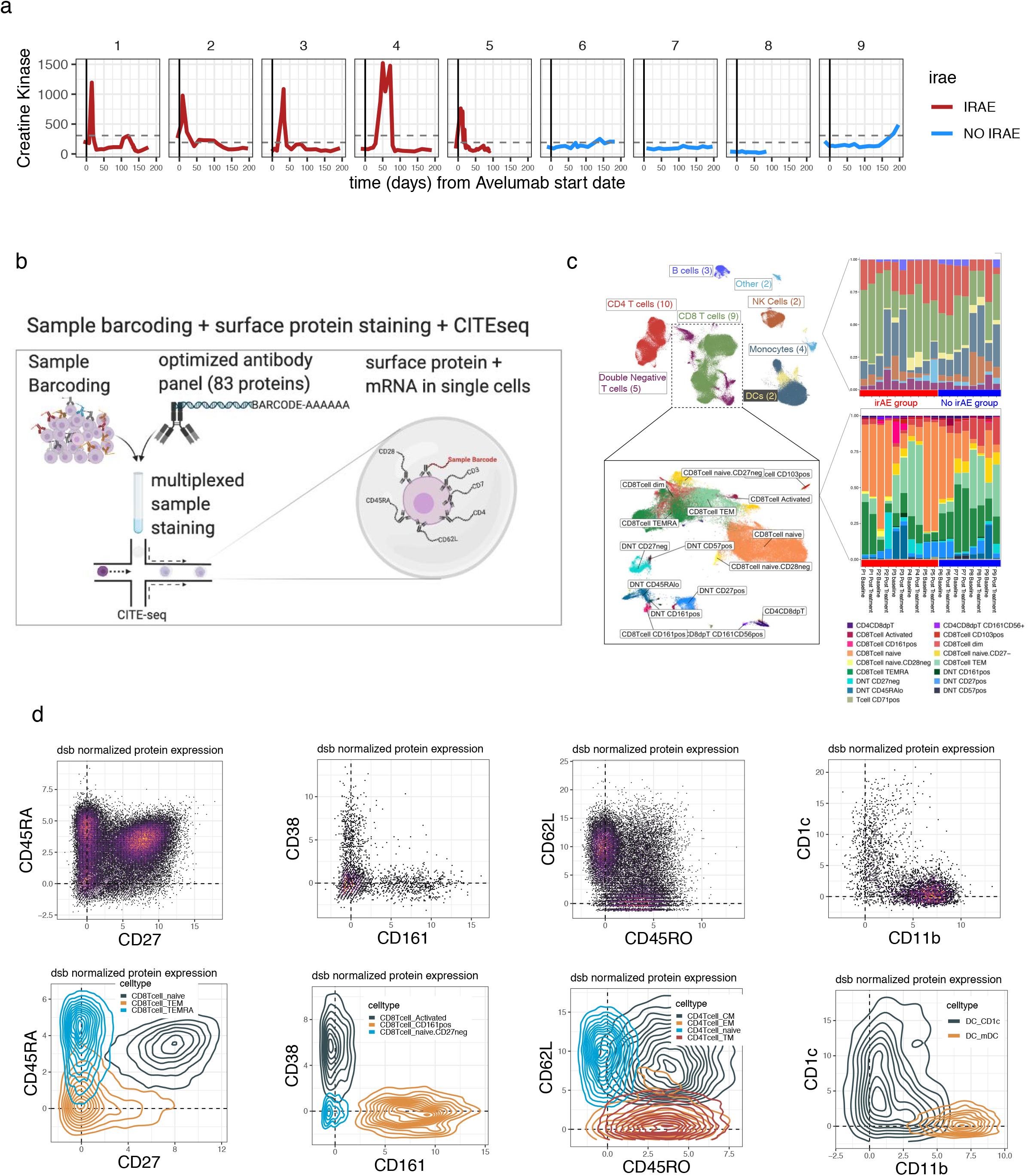
a. Blood creatine kinase (CK) levels (y-axis) vs. time from the initiation of avelumab treatment in patients profiled with CITE-seq b. Sample multiplexing and CITE-seq experiment scheme. c. Left–Uniform manifold approximation projection (UMAP) of PBMC colored by main immune cell lineages with a subset of the map expanded (inset) containing the T cell subsets indicated in the box. Right–the distribution of the number of cells per subset for the clusters shown (top, main lineage, bottom, the T cell subsets shown in the bottom UMAP plot). d. CITE-seq dsb normalized and denoised surface protein expression from single cells in bi-axial plots with the corresponding cells in density plots colored by the spectral clustering annotation. The protein phenotypes align with known canonical cell types which enhances the interpretability of mRNA states associated with the clinical outcomes defined in Fig. 1a.

**Supplemental Figure 2.**
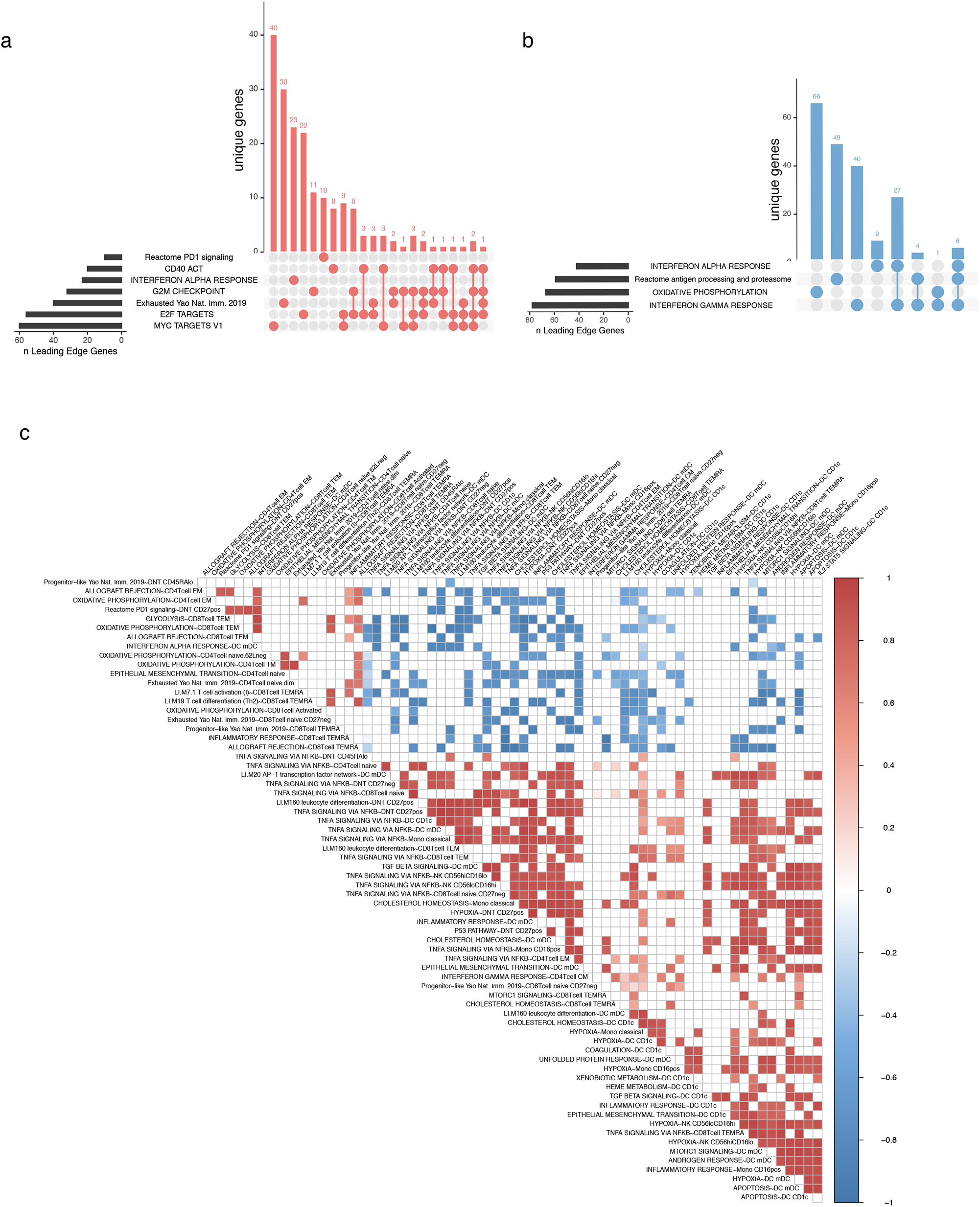
a. UpSet plot of the intersection of leading edge genes for cell state type I (treatment effect) in CD38++ CD8+ T cells shown in Fig 1c. b. As in (a) for classical monocyte enrichments shown in Fig 1e. c. An expanded version of the baseline cell state correlation map shown in fig 2d. Sample level baseline pseudobulk expression correlations of temporally stable baseline states associated with development of later irAE. Each box represents the Pearson correlation coefficient (two sided) of donor pseudobulk data with all correlations with FDR adjusted p value < 0.02 not shown..

**Supplemental Figure 3.**
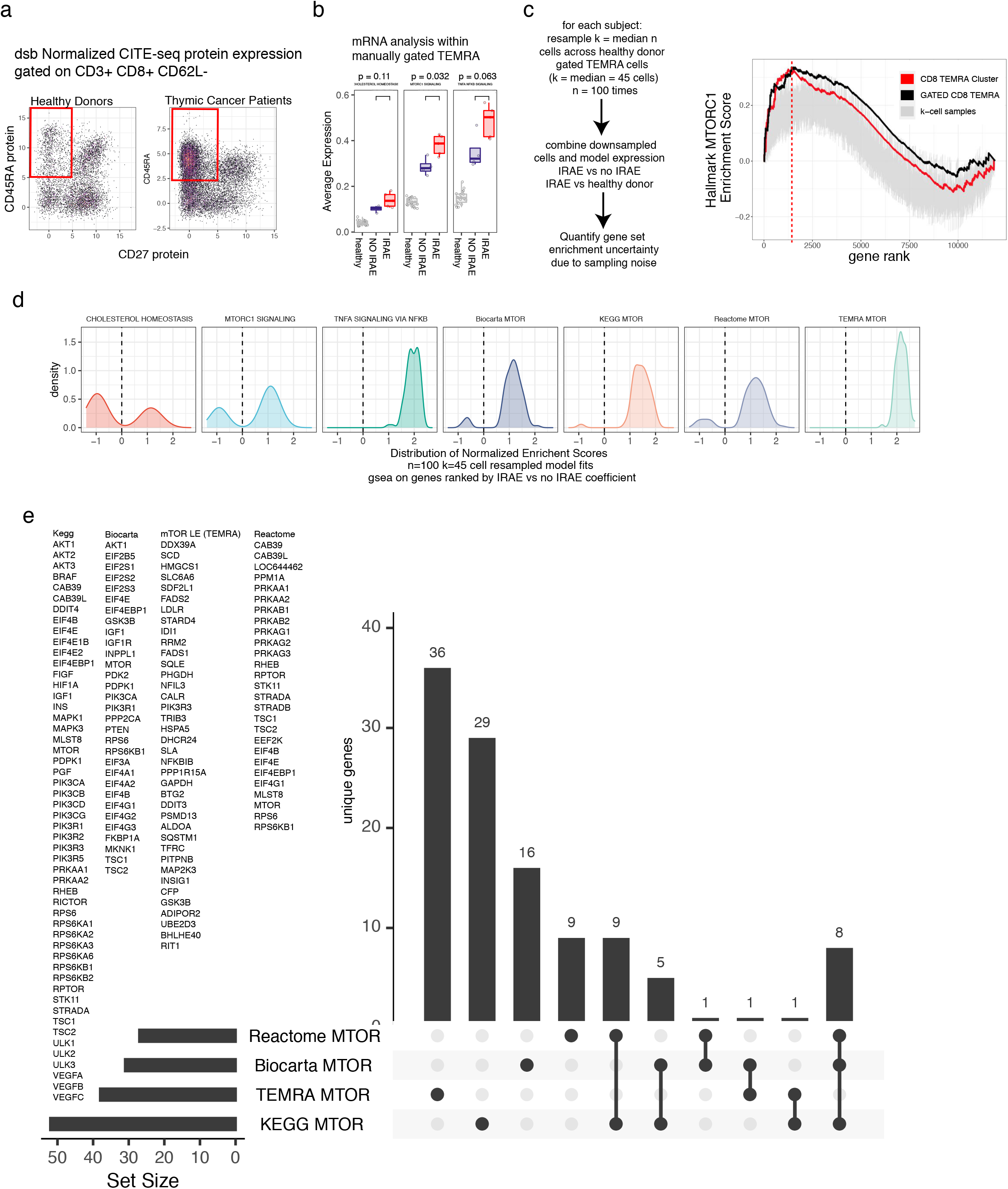
a. Manually gating CD8 TEMRA cells from healthy donors and thymic cancer patients as CD3+ CD8+ CD62L-CD45RA+ CD27-based on dsb normalized protein expression. b. Average expression of the leading mTOR-LE gene signature in the manually gated subsets. c. Robustness assessment of mTOR signature enrichment in irAE vs non-irAE thymic cancer patients and healthy donors; k=45 random cells were re-sampled from each donor to form a downsampled pseudobulk library the resampling procedure was repeated 100 times each followed by a full reanalysis of the weighted pseudobulk model with gene set enrichment based on effect size for the contrast indicated. Grey lines reflect the n=100 down sampled analysis with enrichment from the analysis using the full data (all cells from each donor) shown as the black line and the original enrichment signal from the TEMRA cluster in red. d. As in c; showing the full distribution of normalized enrichment scores across the k-cell permutation testing procedure. e. UpSet plot of mTOR signatures tested in (d) with gene membership in each module shown.

**Supplemental Figure 4.**
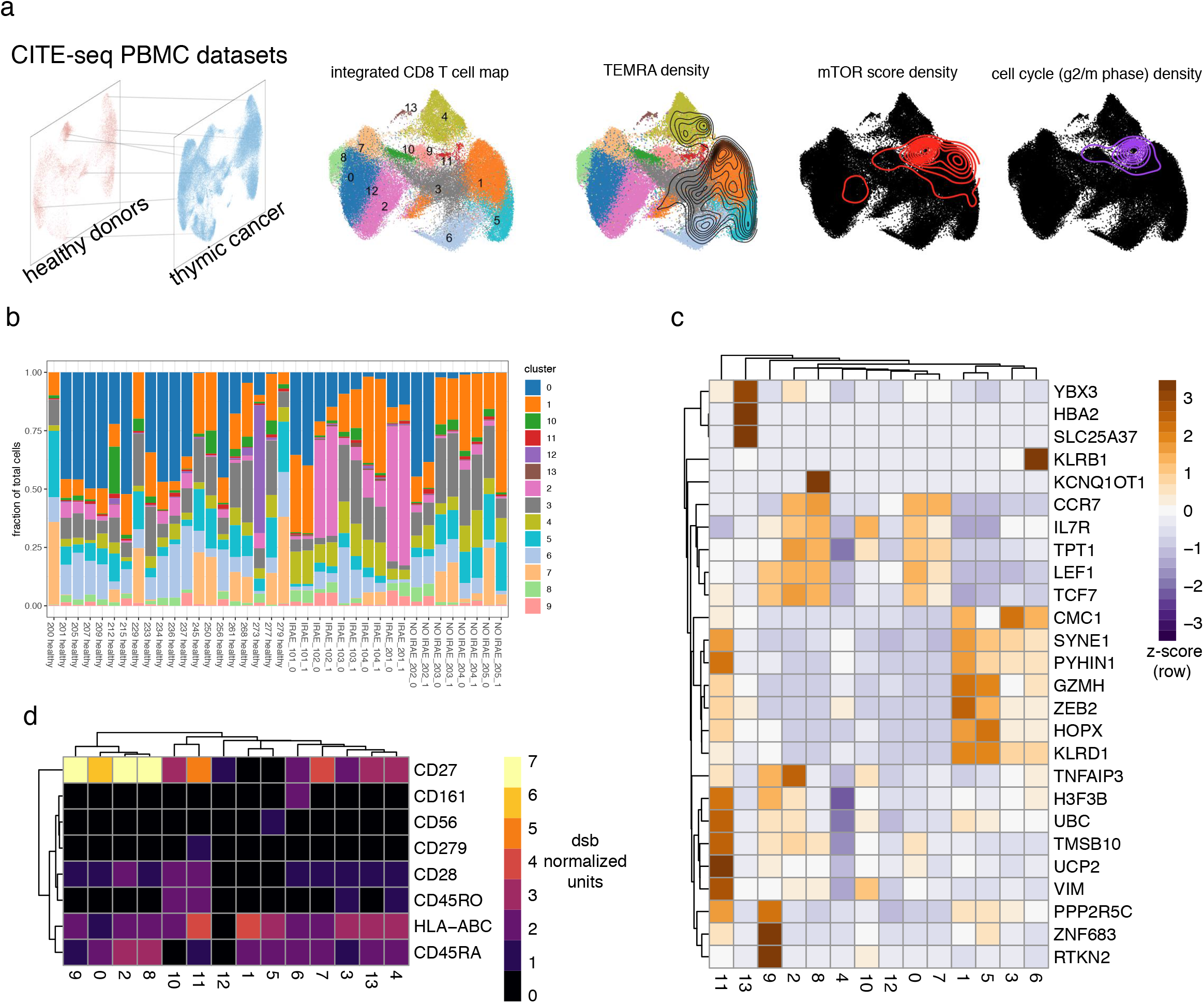
a. CD8 T cells from the n=20 healthy donor cohort were used as a reference dataset on which to project thymic cancer patient cells to form an integrated CITE-seq healthy and cancer T cell map. Cells are colored by integrated assay clustering using Seurat based on 2000 genes regularized Pearson residuals values. The density of the TEMRA cells from the thymic cancer cohort are shown overlaying the integrated mRNA based clusters. The density of cells with mTOR score > 3 absolute deviations from the median mTOR signature score is shown second from right. The right-most UMAP is as above, but for the density of g2m phase score. b. The proportion of cells from each donor belonging to each cluster in the integrated clustering. c. Differential expression of markers between clusters based on regularized pearson residuals with donor effect regressed out, ROC test implemented in Seurat. d. dsb normalized protein expression in clusters as in (c).

**Supplemental Figure 5.**
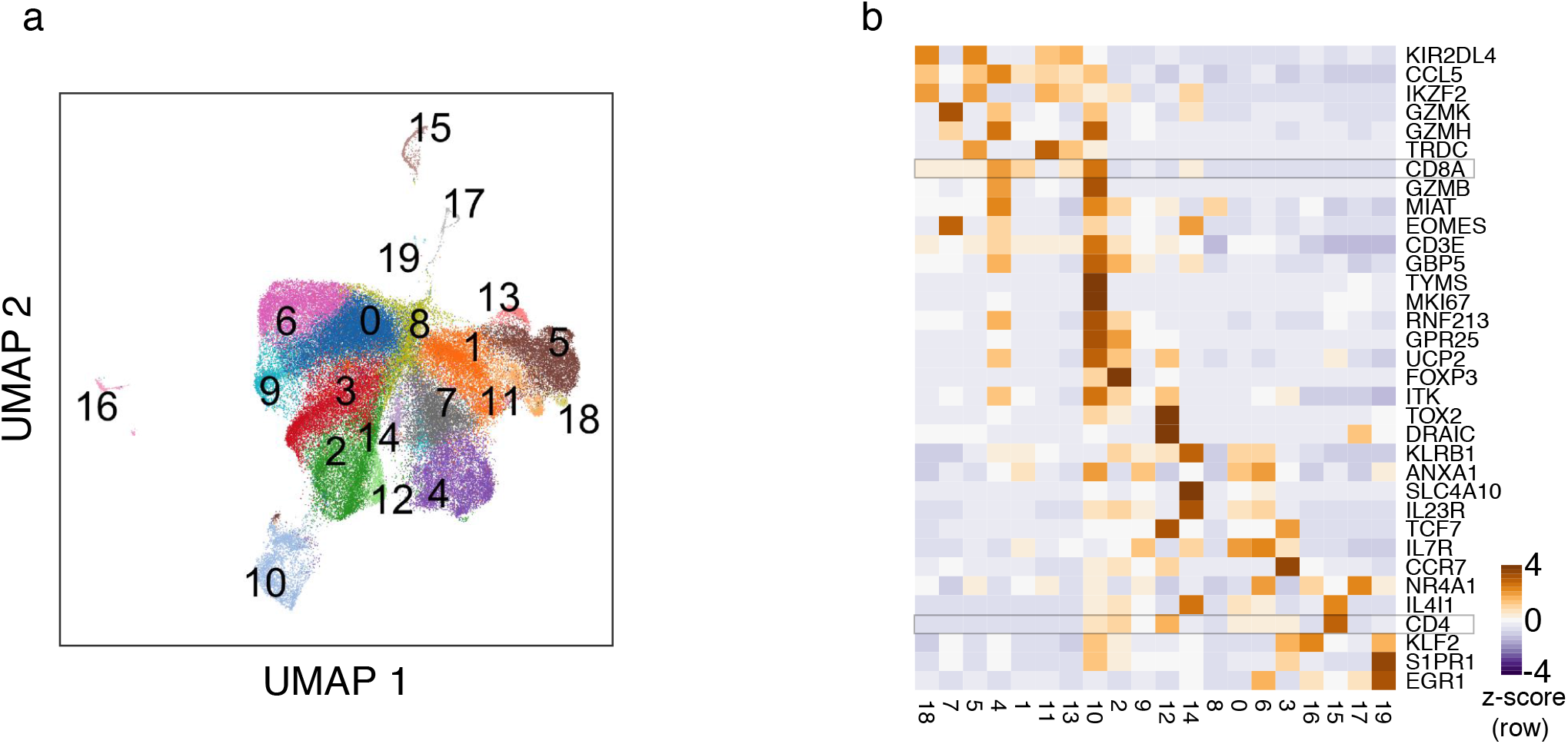
a. UMAP projection of CD3+ cells from melanoma and healthy colonic biopsies; colors indicate clusters derived from graph-based clustering based on 30 principal components from regularized Pearson residuals of 2000 genes. b. Average cluster expression of select genes as shown in Luoma et al.

